# UNDESRTANDING MESENCHYMAL STEM CELL IMMUNE POTENCY: A MORPHOLOMIC AND LIPIDOMIC PERSPECTIVE

**DOI:** 10.1101/2022.05.25.493514

**Authors:** Priyanka Priyadarshani, S’Dravious DeVeaux, Bobby Leitmann, Kejie Rui, Edward A Botchwey, Luke J. Mortensen

## Abstract

Interest in human mesenchymal stem cells (MSCs) as an immune therapy has been on the rise for the past two decades with cutting edge research yielding promising results, but there are currently no MSC therapies approved by the food and drug administration (FDA). Failure of MSCs to translate as a therapy has been reported by the National Cell Manufacturing Consortium (NCMC) to be due to a lack of reliable potency metrics and sufficient understanding of the mechanism of action. Here we show that cell membrane components are a good candidate to interrogate the MSC immunomodulatory mechanism of action and provide a method to increase MSC potency through the sphingolipid pathway. We found that high and low indolamine-2,3-deoxygenase (IDO) potency cells have distinct morphological signatures that is also reflected in the sphingolipid activity, with low IDO potency cell lines having low sphingomyelinase activity and high IDO potency cell lines having high sphingomyelinase activity. Perturbation of the salvage pathway with the addition of exogenous neutral sphingomyelinase not only shifted morphological signatures to a high potency profile, but also significantly increased IDO activity within both high and low IDO potency donors. These results provide a proof of concept for the engineering of MSC immunomodulation and provides further evidence for the role sphingolipids in MSC immunomodulation that can enable further investigation.

## Introduction

Human mesenchymal stromal cells (MSCs) have exhibited great promise to reduce immune and inflammatory responses in vitro and in animal models. This has led to an explosion of clinical trials over the past several years from 142 in 2010 to 895 at the beginning of 2018 (clinicaltrials.gov), but there are currently no FDA approved MSC therapies (FDA.gov). Therefore researchers, industry, and regulators have worked to identify and address challenges to creating high potency cell therapeutics at a relevant manufacturing scale.

Many of the most promising clinical trials for MSCs leverage their immune suppressive activity. MSCs signal immune cells through direct receptor-mediated interactions, vesicle communication and cytokine release mechanisms. The combination of these mechanisms has been suggested to polarize macrophages and dendritic cells [3], induce apoptosis and a decrease in proliferation in T cells [4, 5], and stimulate T-regulatory cell fates [6, 7]. A major barrier to the translation of MSCs in clinical trials is that this complex mechanism of action has made it difficult to identify a unifying underlying biological mechanism that could be leveraged to monitor and enhance cell potency during a manufacturing process.

The physical processes that are associated with these immune-related cell activities (e.g. migration and receptor binding, vesicle release, mitochondrial activation) often have inherent distinct phenotypic outcomes that alter cell shape, or morphology, and can be observed using imaging techniques. Recently, the regulatory networks and principles that govern physical changes in cell morphology definitions have begun to be described and applied to understand temporal perturbations of cell morphological dynamics. Even a static morphological snapshot reflects underlying phenotypic dynamics and is sensitive enough to detect and predict outcomes such as stem cell fate and cell response to drug treatment. For MSCs, morphological phenotype has been associated with their differentiation into osteoblasts and chondrocytes, as well as directly with immune suppressive activity [8, 9]. Klinker and colleagues uncovered MSC morphological signatures that were correlated with MSC immune activation by IFN-gamma stimulus, as well as MSC suppression of CD4+ and CD8+ T-cell activation.

Ensemble measurements of cell imaging phenotypes allow inference of cell state and activity, and so are suggestive of underlying network regulation and homeostasis at the genetic, gene expression, and metabolite level. We focus on a group of lipids known as sphingolipids with structural and signaling functions that play a critical role in regulation of the actin cytoskeleton and cell shape through adhesive linkages, filapodial/lamellipodial protrusions, and endocytosis/exocytosis processes. Regulation of the SL networks are closely linked with inflammatory and immune processes, and the activity of tolerizing enzymes such as IDO. The bioactive signaling lipid S1P alters plasma membrane protrusions in stromal cells through the Rho/Rac migratory axis by acting on the S1P receptors and either stimulating migration through S1PR1 or inhibiting migration through S1PR2. In addition, sphingolipid signaling by ceramide and S1P differentially modulates proteins that link the actin network to the cell membrane known as the ezrin, radixin, and moesin (ERM) proteins thereby influencing cell adhesion, lamellipodia/filopodia formation, and cell migration, which are modulated in part through the activity of sphingomyelinase. Finally, the dramatic cytoskeletal and membrane rearrangements required for endocytic invagination and exocytic vesicle creation are regulated by biophysical and signaling means requiring de novo lipid synthesis, neutral ceramidase activity, and sphingomyelinase activity. When mutations occur that alter the balance of the sphingolipid network, defects in inflammatory and immune regulation often result along with impairment of skeletal tissues [12–16]. The sphingolipid network therefore exists at a unique nexus of lipid signaling, lipid biophysics, and cytoskeletal control; and is reflected in cell morphological properties [10, 11].

In this work, we hypothesized that the sphingolipid network profiles could serve as a biophysical and signaling metabolic link between MSC morphology and immune suppression. We therefore investigated the morphology of 9 MSC donors with varying immune suppressive activity and secretory profiles, and probed the correlation between overall cell shape and regulation of specific lipid-regulated cell compartments. We further examined the sphingolipid profiles of each MSC donor sample and their responses to IFN-gamma immune stimulation. We then identified key lipid modifying enzymes whose activity was correlated enhanced response to IFN stimulation, and used lipid modifying treatments to boost the immune suppressive activity of the cells by stimulating lipid conversion reactions, with a direct readout based on the morphology of the cells. This approach to use imaging to identify underlying metabolic features that can be dynamically tuned to enhance the immune suppressive behavior and improve therapeutic efficacy of culture expanded MSCs.

## Results

### 1. Morphology Signatures Differ Among Different Human MSC Donors

It has been shown that morphological signatures in MSCs are indicative of their function and distinctly separable by morphology after interferon gamma stimulation [8]. We first tested two MSC donors, a high IDO potency donor (lot RB137) and a low IDO potency donor (lot RB49) under interferon gamma stimulation and normal cell culture conditions. Principal component analysis was conducted on morphology data and explained the majority of the variance in the data set (90.8%) with PC1 explaining 70.2% and PC2 explaining 20.6% of the variance in the data. After interferon gamma stimulation, both the high and low potency donor experienced a left shift in morphology along PC1and as expected, the high potency donor’s morphology signature after interferon gamma stimulation was drastically different than the other 3 conditions. Interestingly, we found that under normal cell culture conditions, the high IDO potency donor separated well from the low IDO potency donor along PC1. A look at the PC1 loadings within the loading plot indicates that features that were indicative of cell length and spread (ex: MeanRadius, Max Radius, Extent, FormFactor) were important in the separation seen along PC1. PCA on high and low IDO potency cells at basal conditions shows a clear divide between the two groups along PC1 Figure 1A. Three of the features that most contributed to the separation seen in PC1 (Extent, FormFactor, MinFeretDiameter) were used to create a 3D plot of high and low IDO potency cells using differential features derived from the ratio in morphological signatures between stimulated and unstimulated cells Figure 1B. A clear separation between high and low IDO potency can be seen which further indicates distinct morphological signatures for these two donors. Using PC1 as an indicator of MSC IDO potency, a color-coded meter was created and then cells were false colored based on where they fell within the meter in a given image. We found that high IDO potency cells tended to have a more heterogeneous population as seen in Figure 1C, while low potency MSCs had a more homogenous population of small relatively round cells. We then looked at the density plots to investigate the differences in population distributions of features that exhibited significant differences between distributions seen in Figure 1B using the Kolmogorov–Smirnov test (KS test). For the Area and MaxRadius features, the KS test shows that all of the donors have significantly different population distributions from each other except donors RB139 and RB180, which is reflected in the population distribution plot seen in Figure 1 (D,E). A KS test on the MaxFeretDiameter feature interestingly shows that the RB137 (high potency) and RB49 (low potency) are more similar to each other than they are to the other two intermediate IDO potency donors Figure 1F. Lastly, a KS test on the FormFactor feature shows that all of the donors have significantly different population distributions from one another except RB49 and RB180 (Figure1F. The differences seen between donors based on the population distribution curves using morphology further suggests that each donor has a distinct morphological signature. We then expanded our donor pool to include two intermediate IDO potency MSC donors and performed partial least squares discriminant analysis (PLSDA) on all four donors to test the predictive power of MSC morphology on IDO potency. We found the PLSDA had a cross-validation misclassification error was 2.5% for predicting the IDO potency of the four donors and that each of the donors seemed to separate along the LD2 axis by IDO potency. While RB137 and RB49 (high vs low IDO potency) were clearly separable along LD2, the two intermediate donors (RB180 and RB139) were morphologically similar (Figure 1H).

**Figure 1.**
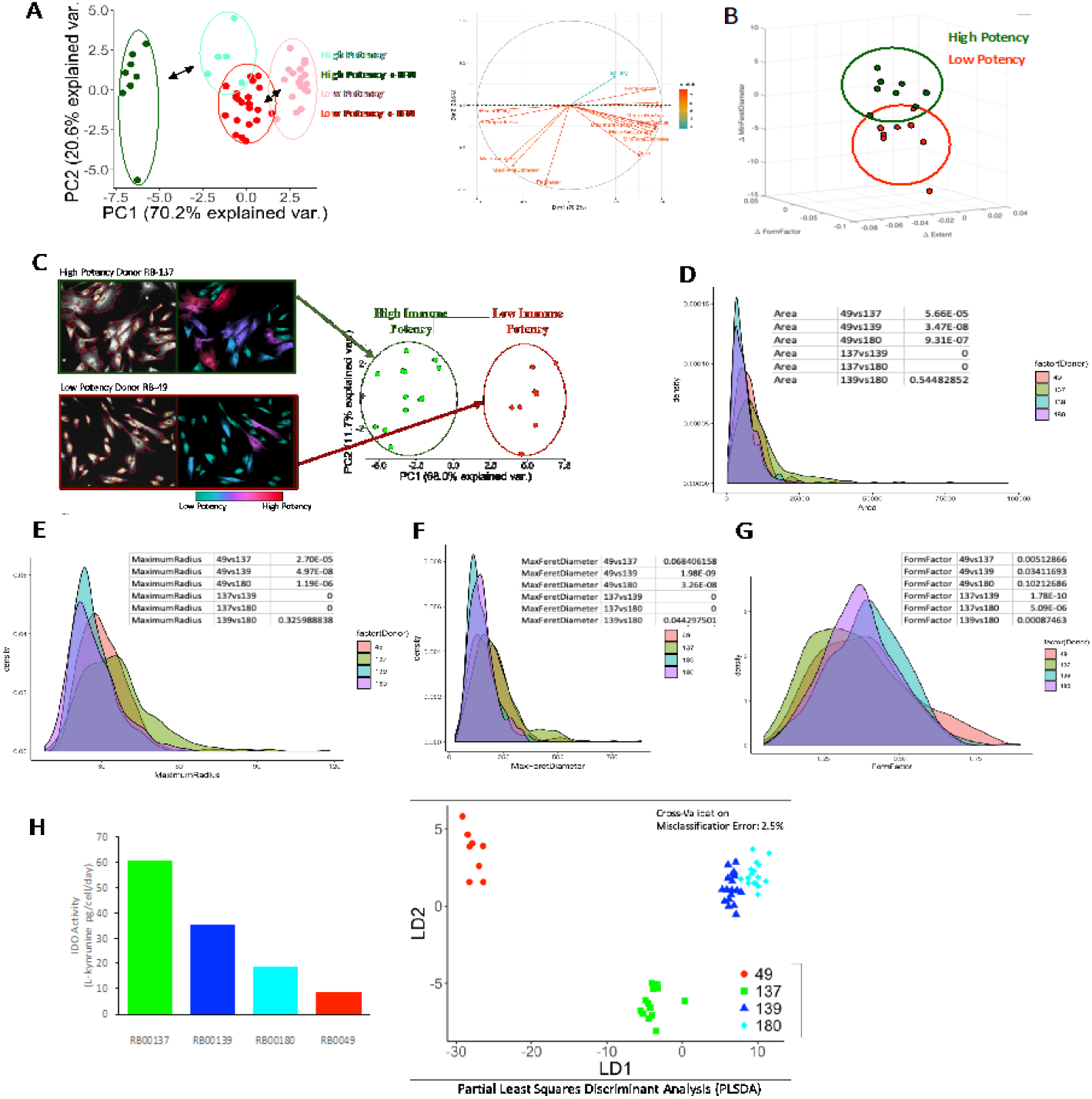
Characterization of morphological features show predictive potential of MSC immunomodulation. A) morphological features of high and low IDO potency MSC donors under normal MSC culture conditions were obtained with CellProfiler and analyzed with PCA. Morphological supercell features clearly separate high and low potency MSCs along PC1 indicating distinct morphological differences between high and low potency donors. B) Stimulation of MSCs with IFN shows a left shift along PC1 for both high and low potency donors with the high potency donor having a larger morphological signature change after stimulation than the low potency donor. C) Delta features (feature ratios of stimulated/unstimulated) selected from highly weighted features in the PC loadings (left) plotted onto a 3D plot shows that measurements in the PC loadings yield separation between high and low IDO potency donors. D) The addition of two intermediate IDO potency (RB00139 and RB00180) donors in PCA shows that their morphological signatures lie somewhere between the high and low potency cell lines. E) A look at the distribution plots of the donors shows that these four donors have differences in the distribution of morphology in the donor cell population. A ks test confirms that there is significant evidence that donor 137 and 49 have significantly different distributions from each other and the other two intermediate donors while donor 139 and 180 are fairly similar to one another. H) PLSDA on morphological features shows separation along LD2 by donor potency with a 2.5% misclassification error with the cross-validation method.

### 2. Lipidomic Profiles Reveal Sphingolipid Activity Differences in High versus Low IDO Potency Donor Groups

For lipidomic analysis MSC donors were split into two groups (high vs low IDO potency) using the mean of the group as the threshold as seen in Figure 2A. Lipidomic analysis was conducted using high performance liquid chromatography tandem mass spectrometry (HPLC-MS/MS) to profile eight different donors. In Figure 2B a Pearson correlation heatmap was created to determine differences in lipidomic metabolism between high and low potency donors. We found that there were several key differences between the two groups. The most outstanding being the relationship between ceramides (Cer) and sphingomyelins (SM). In the high IDO potency group there was a negative correlation between high and low potency cells indicating that there is high conversion of one metabolite to another. In low IDO potency cells there was a positive correlation between SM and Cer indicating that there is little conversion between these two sphingolipids. A look at long chain Cer and SM levels after stimulation of high and low IDO potency groups with interferon gamma shows no significant difference between stimulated and unstimulated groups. On the other hand, under normal cell culture conditions there were significant differences between low and high IDO potency in all four sphingolipids. Levels of long chain Cer and SM were consistently lower in the high IDO potency group when compared to the low IDO potency group as seen in Figure 2C. C24:0 SM had the most significant difference between high and low IDO potency groups and a further look at the levels of the downstream sphingolipid C24:0 Cer shows no difference between low and high IDO potency groups, but C24 SM levels were significantly different between high and low potency groups indicating that high IDO potency groups metabolize SM more than low IDO potency groups (Figure 2D). A further look at the ratio between Cer and SM showed that the C24:0 Cer/SM ratio of and the C24:1 Cer/SM ration were significantly different between high and low potency groups further indicating that the conversion of the long chain and complex long chain SM to Cer is much higher in high IDO potency cells vs low IDO potency cells (Figure 2E). An NBD tracer assay tracking the conversion of sphingomyelin to ceramide indicating sphingomyelinase activity showed that sphingomyelinase activity in high IDO potency cells is significantly higher than low IDO potency cells (Figure 2F) and there is conversion of sphingomyelin to ceramide in high IDO cells (Figure 2G).

**Figure 2.**
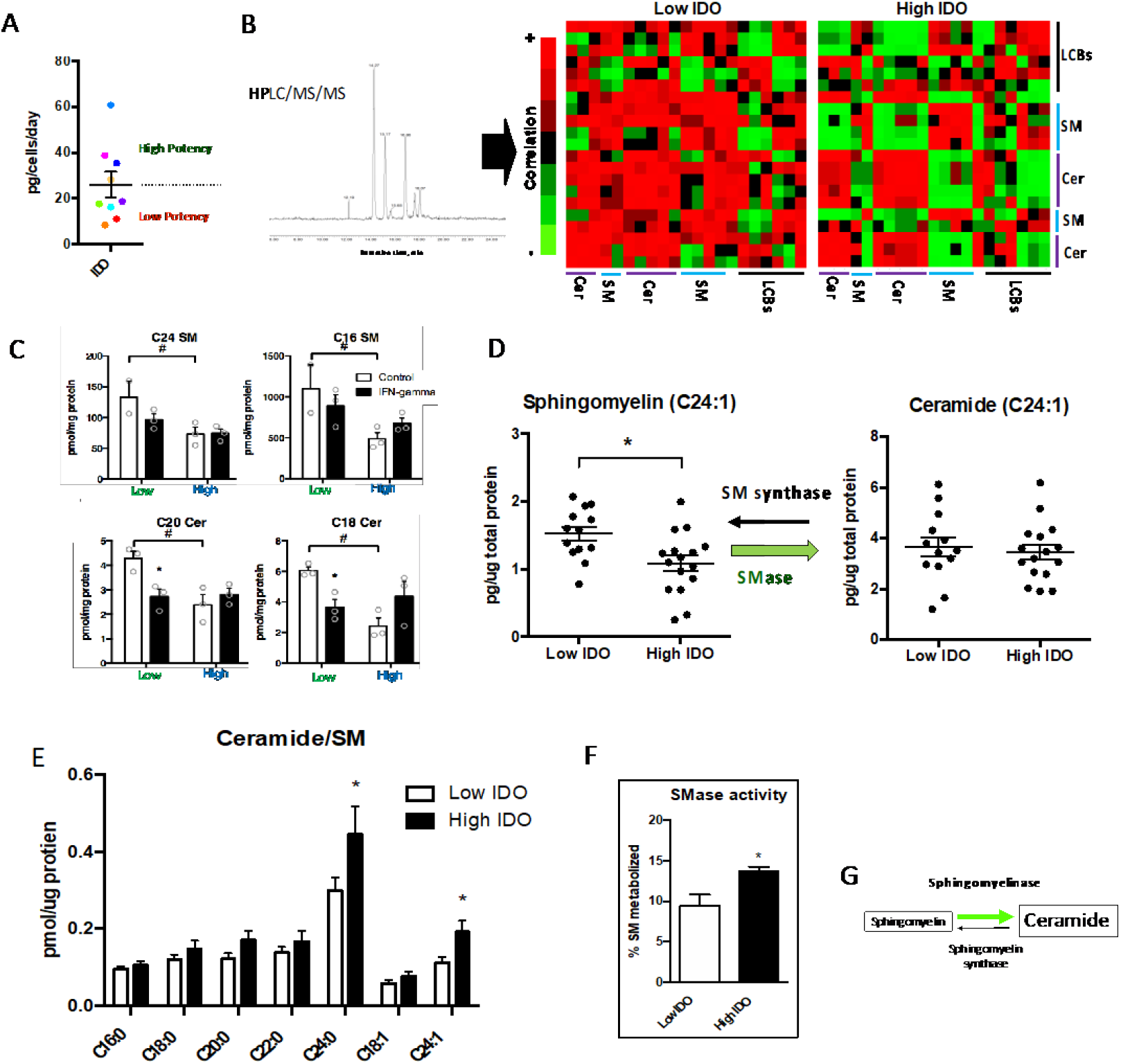
A deeper look into membrane components with lipidomic analysis shows sphingomyelinase as a key underlying factor that distinguishes high and low IDO potency donors. A) Donors were grouped into high and low IDO potency with the threshold for high vs low IDO potency being the mean of the group. B) Pearson’s correlation heatmap of the lipid data shows distinct differences between a high and low potency donor. C) Level of long chain ceramides in low and high potency cells. D) Plot showing level of C24:1 in low and high potency cells. E)Plot showing major of different cermide in low and high potency cell line. F) Examination of the sphingomyelinase activity shows that high IDO cell lines have higher neutral sphingomyelinase activity. G) Conversion of sphingomyelin to ceramide in high IDO cells.

### 3. Perturbation of Sphingolipid Pathway with Sphingomyelinase Shifts Morphology and Increases MSC IDO Potency

Sphingolipids have important roles in both MSC immunomodulation and morphology phenotype Figure 3A. To test the effects of sphingomyelinase activity on MSC morphology cells were treated with sphingomyelinase and morphological features were extracted and compared to cells that had no treatment or had a sphingomyelinase inhibitor (GW3869). Morphological signatures in PLSDA show that cell stimulated with sphingomyelinase have a distinct shift in morphology phenotype when compared to control as seen in Figure 3B. Furthermore, we tested the effect of sphingomyelinase + interferon gamma treatment on IDO activity and found that IDO activity significantly increased in both high and low IDO potency donors (Figure 3C).

**Figure 3.**
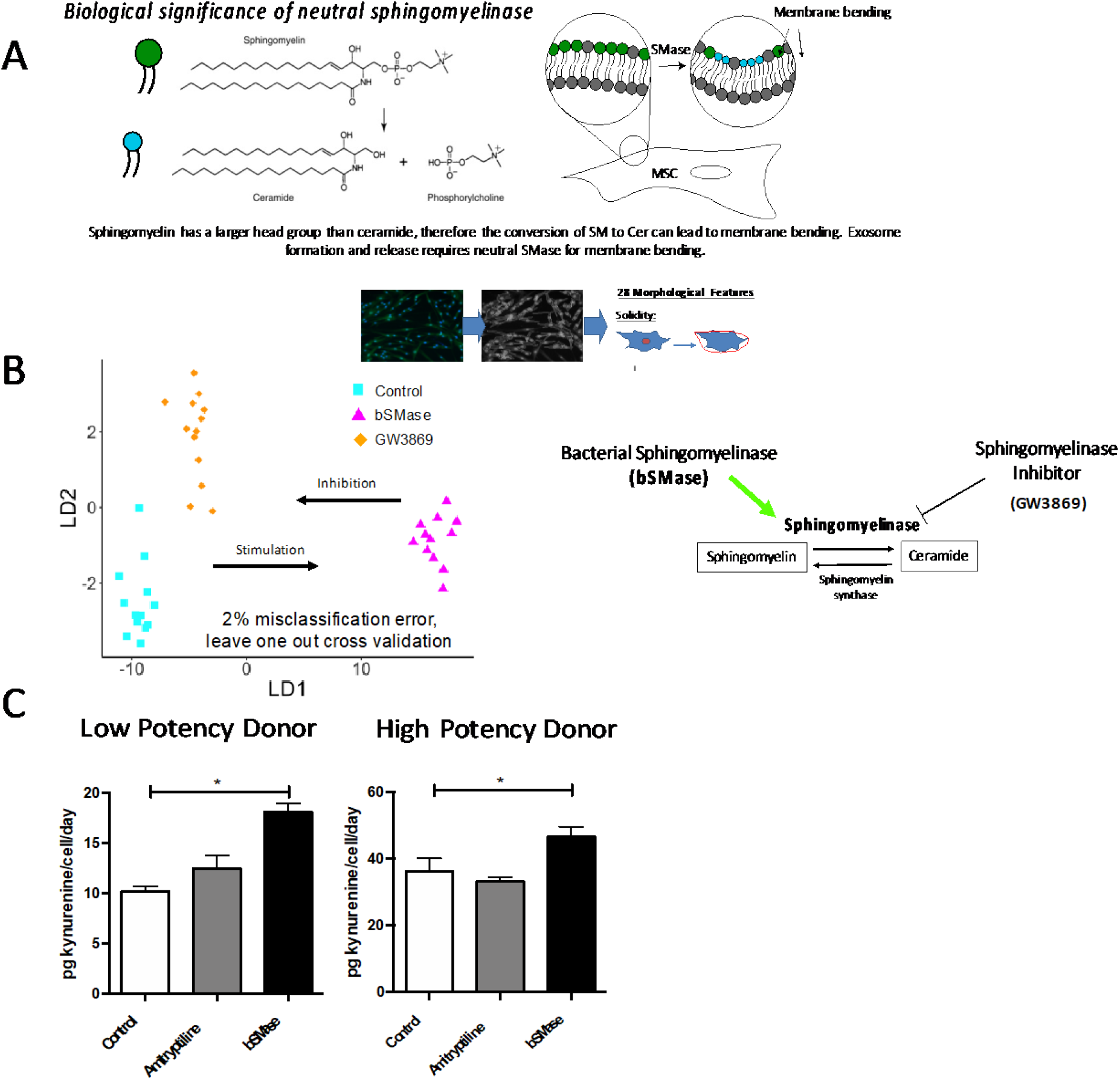
Sphingolipids are important components of the cell membrane and ratios of ceramide and sphingomyelin cause membrane bending leading to morphological changes in the cell (A). Supplementation of bacterial derived neutral sphingomyelinase to increase ceramide production creates distinct morphological features (B). There is a shift in morphology phenotype after treatment with Neutral SMase when compared to control. Inhibition of SMase activity creates a shift back to a morphology phenotype that is similar to control. Supplementation of MSCs with bacterial derived neutral sphingomyelinase increases IDO activity in both high and low potency donors (C).

### 4. Treatment with bSMase increases secretion of extracellular vesicles

Since we know that treatment with bSMase enhance the conversion of sphingomyelin to ceramide in high IDO cells, we further wanted to determine how this conversion help in functional properties of the cells. We treated the cells with bSMase and performed longitudinal imaging. The images show increase in secretion of small particles in bSMase treated cells (Figure 4A). As IFN-ɣ is shown to enhance the functional properties of MSCs, we treated the cells with combination of IFN-ɣ + bSMase and compared the release of extracellular vesicles (EVs) with other treatments like only IFN-ɣ, IFN-ɣ +GW and control. The release of EVs is shown be higher in IFN-ɣ + bSMase group compared to all other groups (Figure 1B). Moreover, there is also increased concentration small particles in IFN-ɣ + bSMase group compared to the only IFN-ɣ group (Figure 1C). To further demonstrate the presence of EVs, we stained the cells with CD63. The images stained showed higher concentration of CD63 in bSMAse treated vs untreated group (Figure 4D).

**Figure 4.**
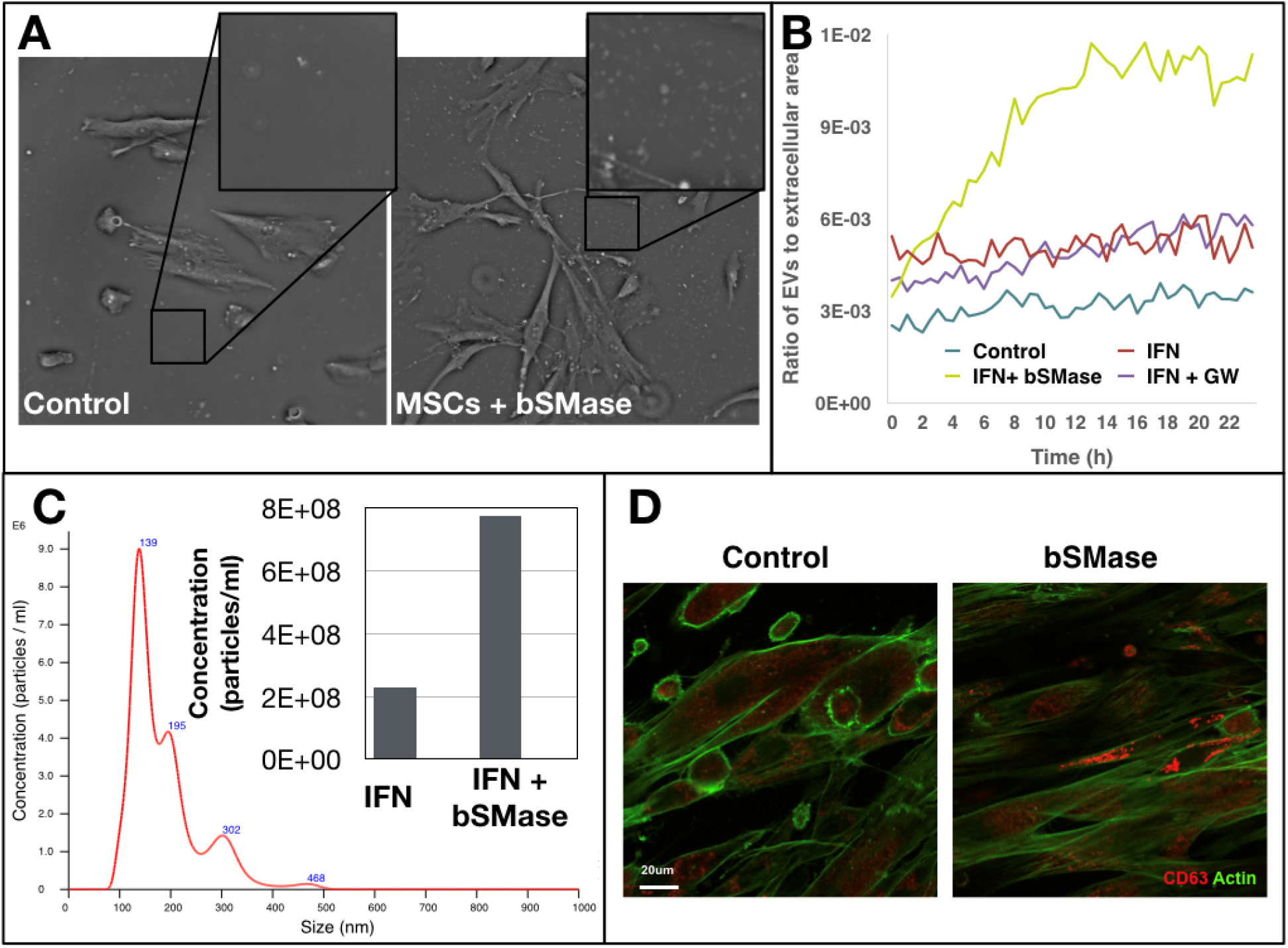
Longitudinal phase imaging of MSCs shows increased number of small particles after stimulation with IFN-ɣ + bacterial sphingomyelinase (bSMase), which was lacking with the addition of the SMase inhibitor GW(A). When these particles were quantified using a NanoSight, a size distribution in 100-300 nm range was detected (B). With an increase in the total number of extacellular vesicles (C inset). Ceramide-rich extracellular vesicles are thought to be created inside the cell in multivesicular bodies that are labeled by CD63, which appeared to be increased with the addition of bSMase (D)

## 3.4 Materials and Methods

### Cell Culture

Nine different donors of human bone-marrow derived MSCs were obtained from RoosterBio Inc. Cells were expanded once in RoosterBio expansion media in standard growth conditions (in the dark at 5% CO2 and 37C with humidity) and frozen down for cell bank. Depending on conditions to be tested cells were either experimented on in RoosterBio media or standard hMSC media: α-MEM, 10% FBS, 1% L-glutamate, 0.1% penicillin-streptomycin.

### Morphological Assessment

Cells were seeded at 3,000 cells/cm2, allowed to adhere for 24hrs and then treated with various treatments for an additional 24hrs then fixed with 4% (w/v) paraformaldehyde for 15 minutes and then rinsed 3 times with PBS. Fixed cells were then stained with 300nM DAPI in 0.1% Tween20 for 10 minutes and 20uM FITC for 30 minutes. Images were captured with VTI Array Scan High Content Imager (ThermoFisher). Images were then processed using CellProfiler and 28 features were extracted from the images: 14 nuclear features and 14 cell body features.

### Lipidomic Analysis

Cells were harvested at 80% confluence and frozen in −80C freezer at a concentration of 1 million cells/350uL in 400uL of PBS. Lipidomic profiles were obtained using High Performance Liquid Chromatography (HPLC) conducted by

### IDO Activity Assay

IDO activity assay was performed as previously described in Daubner et al 1993 with some minor changes. Cells were incubated with IFN-*Y* for 24hrs then 100uL conditioned media was collected in round bottom 96-well plate and 50uL of 30% (w/v) trichloroacetic acid (TCA) was added to each sample to convert N-formyl-kynurenine to it’s detectable form L-kynurenine. Samples were then centrifuged at 1000g for 5 minutes. 75uL of supernatant was transferred to a flat bottom 96-well plate and 75uL of 2% (w/v) 4-(dimethylamino)benzaldehyde in acetic acid (Ehrlich’s reagent) was added to each sample. Absorbance was read at 490nm on a plate reader. Data were normalized by media volume in culture, cell number, and days in culture to get the units of pg kynurenine/cell/day. Standards were done in triplicate and in respective medias using L-kynurenine supplement. Cell number was obtained by PicoGreen quantification. Fixed adherent cells were incubated for 30 minutes in PicoGreen and then immediately read on a plate reader at 480nm excitation and 520nm emission as described by ThermoFisher’s protocol. Serial dilution 1:2 of cells from 50,000 cells to 3,125 cells was performed for standard curve for each donor line.

### Statistical Multivariate Analysis

All multivariate statistics methods were performed using RStudio. Morphology feature data was first converted to supercell format by averaging the morphological features of a random set of 30 cells within a condition. Supercell data was then run through principal component analysis using the correlation matrix because feature scales were drastically different. Predictive models using morphological data were created using partial least squares discriminant analysis.

## Discussion

Indolamine-2,3-deoxygenase (IDO) is widely known for its anti-inflammatory action on multiple immune cell types Human MSC’s are known to increase IDO activity upon stimulation of IFN and its rate of activity has been shown to be predictive of overall MSC immune suppression on peripheral blood mononuclear cells (PBMC) by down regulating PBMC proliferation and activation markers [4]. With its ease of use, the IDO activity assay is a good candidate for validation of MSC function in the development of fast screening potency metrics and validation of possible alterations of potency. In this research, we were able to show that manipulating aspects of the cell membrane has functional outcomes for MSC immune suppression. We found that different IDO potency donors had unique morphological signatures that can be used for potency prediction. We also found that sphingolipid profiles were different between high and low potency donors leading to a potential path for potency alteration. A deeper look at these sphingolipid networks led us to identify that the conversion of long chain and complex sphingomyelin to ceramide play key roles in high vs low IDO potency cells and provided an avenue for potency alteration of MSCs. From this, we tested the effect of neutral sphingomyelinase, an enzyme responsible for the conversion of long chain and complex sphingomyelin to ceramide as seen in Figure XX, on MSC morphology and IDO activity. We found that sphingomyelinase + IFN treatment created a unique morphological signature and that IDO activity of both high and low potency donors increased after treatment.

MSC immunomodulatory properties have been established since the late 90s [17], but several decades later we still know little about the mechanism of action for MSC immunomodulation [18]. IDO activity in MSCs is a promising functional measure of immune potency in vitro, because its activity has been shown to be predictive of overall MSC immune suppression of activated PBMCs [4]. A couple of groups found that MSCs significantly lost the ability to effectively suppress PBMC activation after IDO activity was inhibited with the IDO specific inhibitor 1-methyl-tryptophan (1MT) [4, 5]. The IDO activity assay is an easily reproducible functional assay that directly tests MSC activity that has a strong relationship to MSC immune suppression outcome. Its main limitation is that it does not directly test the outcome of MSC function on immune cells such as peripheral blood mononuclear cells (PBMCs). To remedy this, future work will include validation with a PBMC activation assay. The PBMC activation assay, however, has its own limitations such as reproducibility, which can be attributed to the use of multiple donor types from PBMCs and MSCs and their unique responses to each other and variation based on culture conditions and dosing. Validation of one assay with the other can mediate these short comings. We first characterized the IDO activity of each of our donors and found that MSC IDO activity was relatively stable in early population doubling (or passage) donors when compared to reported values by the manufacturer (RoosterBio Inc.). A wide range of IDO potency donors allowed for high and low potency grouping with the mean value of the group being the threshold for high and low potency. Donors were then chosen for further studies in morphology and lipidomic profiles based on their IDO potency to interrogate the morphological phenotype we see.

Morphological phenotypes can reveal a lot about cell function including apoptosis[19], cell motility [20], paracrine signaling and contact mediated cell-to-cell communication [21, 22], mitosis, adhesion [23], and function [24]. Recently, morphological signatures of MSCs after interferon gamma stimulation have been shown to have predictive power of both MSC differentiation potential and immunomodulatory potency [8, 25]. Klinker and colleagues found that a measure of potency could be derived from MSC morphological signatures to discriminate between high and low potency MSC donors [8]. We hypothesized that there are underlying mechanisms behind these morphological signatures that can provide further insight into the mechanism of action of MSC immune suppression, but first we established morphological profiles for our own set of donors that allowed discrimination of IDO potency. We started with a high and low IDO potency donor under interferon gamma stimulation. In Figure 3.2A we saw that principal component analysis on IFN stimulated morphological features yields morphological signatures that are easily separable in the PCA space along PC1, which we expected because Klinker and colleagues found that IFN stimulated morphological features better separate low and high potency MSC donors [8, 9]. We also saw that IFN stimulation caused a distinct shift within the PC space for both high and low potency donors in the same direction (Figure 3.2A). This phenomenon also held true for Klinker and colleagues when they treated human bone marrow derived MSCs with interferon gamma and ran PCA on MSC morphological features [8]. A further look the change in morphological phenotype by looking at the difference in cell shape features before and after interferon gamma stimulation (Figure 3.2B) showed adequate separation between a high and low IDO potency donor along three highly influential features (MinFeretDiameter, FormFactor, and Extent) in PC1 of Figure 3.2B. Similar features were also used by Klinker and colleagues on the same cell lines and saw that donors of different immune potency separated well [8]. MSC immunomodulation is dependent upon stimulation by proinflammatory cytokines such as interferon gamma [26] and interferon gamma stimulation occurs through the JAK pathway leading to phosphorylation of STAT1 and subsequent translocation to the nucleus for transcription genes associated with interferon gamma that affects the overall function of the cell depending on the cell type [27]. STAT1 dephosphorylation on the other hand has been thought to be mediated by adhesion and cell shape through proteins in the actin cytoskeleton and cell membrane [27]. This relationship between IFN and cell shape through the JAK/STAT1 pathway could explain the shift in MSC morphology we have seen after IFN stimulation that is consistent in all MSC IDO potency donors and has also been noted by Klinker and colleagues.

Interestingly we found that MSC morphological profiles of a high and low IDO potency donor at normal growth conditions were also separable along PC1, though not as distinctly as when they are stimulated with interferon gamma as seen in Figure 3.2A. PCA on morphological features of just the high and low potency donor under normal culture conditions in Figure 3.2C further shows a clear separation along PC1, further suggesting that high and low IDO potency donors could be identified with morphology without stimulation. Though this phenomenon has not yet been reported in the literature to be predictive, differences in morphology at growth conditions between MSCs from different donors [28] and tissue [29] sources has been reported and it is widely known that MSC potency varies depending on the multiple factors including donor and tissue source [8, 29, 30]. Expansion of these findings into large donor pool could lead to the identification of MSC potency levels under growth conditions, which would be advantageous from a cell manufacturing perspective, because it would allow for a direct, in line, and fast screen of MSC function without alteration of the cell product before use in a patient. It is well known that MSCs in culture exhibit a heterogeneous population that is easily identifiable through imaging [8, 9, 31, 32], but the relative function of each subtype is not known. Since PC1 explained a majority of the variance in the data (68%) and showed a clear separation between high and low IDO potency, we then used it as a potency measure that could be then be superimposed on images and identify possible subpopulations within a given donor that could be indicative of immune function (Figure 3.2C). We found that there was a difference in cell population distributions between high and low potency cells. Low potency cells had a high number of small and relatively round cells seen in cyan and light blue in Figure 3.2C while high IDO potency cells had more cells that were intermediate in size seen in purple, pink and blue. Identifying the possible functions of these sub populations might reveal an important factor of MSC immune suppression. These subpopulations have been recently investigated by Marklein and colleagues. They found that overall MSC immunomodulation could be predicted by the prevalence of a specific morphological subtypes (relatively medium sized cells) that they dubbed SPy9 [9]. Isolation and selection of such a subtype could lead to less variability in MSC immunomodulation.

We then wanted to look at how population distributions differed between four donors of varying IDO potency based on morphological features. A Kolmogorov–Smirnov test (KS test) revealed that donor population distributions were significantly different from one another with the following variables having the biggest difference between: Area, MaximumRadius, MaxFeretDiameter, and FormFactor (Figure 3.2D). This data suggests that within our cell lines we have varying distributions of morphological phenotypes. This has been studied recently in the aforementioned study that showed stimulation of MSCs with interferon gamma shifted the population distribution of MSCs and shifts occurred most notably in features such as Area, MajorAxisLength (similar to MaximumRadius and MaxFeretDiamater), and FormFactor [9]. From this and our data it seems that size, shape and spread of MSCs appear to reveal functional outcomes.

A clear mechanism of action for the morphological phenotypes seen between different IDO potency MSC donors is not clear, but a key subset of lipids known as sphingolipids are well established to be important in maintaining the cell cytoskeleton and having influence over cell shape [10, 11, 33]. There is also evidence that sphingolipids play important roles in helping regulate immune responses in elements of the innate immune system, cell survival, anti-inflammatory activity, and homing to sites of inflammation/injury [13, 34–36]. This evidence led us to hypothesize that an understanding of the sphingolipid activity within mesenchymal stem cells could elucidate a possible mechanism of action that explains the morphological changes that are seen between donors of different immunomodulatory potency. We ran a pearson correlation heatmap on 22 sphingolipids within four high IDO potency donors and five low potency donors under normal growth conditions to explore the difference in lipid relationships (Figure 3.3B). We found that high IDO potency donors had a strong negative correlation between sphingomyelin and ceramide while low IDO potency donors had a strong positive relationship between these two types of sphingolipids (Figure 3.3B). A similar study in 2016 looked at the lipidomic profile of MSCs after stimulation with TNF-alpha and INF and found that sphingomyelin levels were increased after MSC stimulation, but the role that sphingomyelin played in MSC function, it’s specific activity remained unclear and possible differences between different donors was unclear [37].

We decided to then dig a little deeper to parse out the drastic differences seen between high versus low potency MSC donors in relation to ceramide and sphingomyelin. To see if more ceramide is being produced or more sphingomyelin, we looked at the relative concentrations of different length ceramides and sphingomyelins in low and high potency donors (Figure 3.3C&D). We found that long chain (C24.0) and complex (C24.1) sphingomyelin concentrations were significantly decreased in high potency donors when compared to low potency donors indicating that the negative correlation between the two sphingolipids was towards the conversion of sphingomyelin to ceramide (Figure 3.3C and D). A look at the ratios between same chain length ceramides and sphingomyelins showed that long chain (C24.0) and complex (C24.1) Cer/SM ratio was significantly different between high and low potency IDO donors (Figure 3.3E). This suggests that long chain and complex Ceramides may have specific functions in MSCs related to morphology and immunomodulation. This corroborates a new paradigm in sphingolipid metabolism that states that specific ceramide species and their metabolites have specific bioactive functions in the cell [36, 38].

We then tested the activity of neutral sphingomyelinase, the main enzyme important in the conversion of long chain sphingomyelins to ceramides, and found that the activity was significantly higher in a high potency donor when compared to a low potency donor (Figure 3.3F). The role of sphingomyelinase in the regulation of cell adhesion has been shown in human monocytes where induced sphingomyelinase activity reduces the binding of leukocytes to monocytes with a suggested mechanism of action through the enhanced interaction of lymphocyte function associated antigen −1 with the surrounding cortical actin [39]. The disruption of cell binding by sphingomyelinase through cortical actin has implications on its impact in cell morphology, because sphingolipids have been shown to directly influence the actin cytoskeleton which controls cell shape and function [11, 33].

We then wanted to investigate the relationship between morphology and sphingolipid activity, so we treated one of our donors with sphingomyelinase and investigated the morphological shifts that occurred. Partial least squares discriminant analysis was used to discriminate between the different treatment groups based on morphological feature inputs. We found that the morphological signatures generally split by treatment group along LD1. Comparison of control MSCs with sphingolipid altering factors showed that the addition of a ceramide production inhibitors through the denovo pathway (not shown) and the salvage pathway (sphingomyelin to ceramide) generated similar morphological signatures indicating that the reduction of ceramide from either pathway has similar effects on the overall morphological phenotype. We expected that this would be the case, because the key difference between high and low potency donors was the production of ceramide. Interestingly it has been found that the inhibition of ceramide production through the denovo pathway yielded an altered morphological phenotype in fibroblasts [11]. They found that treatment of Swiss 3T3 fibroblasts with fumonisin B1 (FB_1_), which inhibits the denovo pathway by blocking ceramide synthase conversion of sphingosine to ceramide, affected overall actin cytoskeleton production and maintenance. This resulted in drastically different morphological phenotypes between untreated and treated cells, which was evident in the loss of leading edge lamella, a phenomenon maintained and controlled by the actin cytoskeleton [20], in the treated group [11]. When we added neutral sphingomyelinase, we found that the morphological phenotype was drastically different than the control and inhibition groups as seen in the PLSDA plot in Figure 3.4B. We expected that the addition of neutral sphingomyelinase in a low potency donor would result in a morphological phenotype that is similar to a high potency donor, creating a large shift in morphological signature.

Noting the significantly increased activity of neutral sphingomyelinase in high potency vs low potency MSCs, we hypothesized that exogenous treatment of cells with sphingomyelinase may increase IDO immune potency in both high and low potency donors, so we aimed to investigate the functional outcome of MSCs after treatment with neutral sphingomyelinase. To test this, three different treatments were tested for MSC IDO activity: 1) interferon gamma as our control because activation of MSCs is crucial in the induction of IDO activity, 2) interferon gamma treatment + amitriptyline because inhibition of ceramide production would provide a negative control, and 3) interferon gamma + neutral sphingomyelinase to evaluate the effects on immunomodulatory function. Treatment of cells with amitriptyline did not significantly change the IDO activity of the cells as seen in Figure 3.4 C. This is not what we expected to see and we suspect that compensatory activity of ceramide production may be at play through the other three metabolic pathways for ceramide production which has been seen in the inhibition of the denovo pathway through FB1 on murine hepatocytes and subsequent increase in acid sphingomyelinase activity to compensate for the loss of function [40]. A double inhibition of both the salvage and denovo pathway may be needed to effectively reduce ceramide production to levels that may affect overall IDO immune potency. On the other hand, we found that the interferon gamma + neutral sphingomyelinase treatment groups in both high and low IDO potency MSC donors had significantly more IDO activity when compared to the interferon gamma control (Figure 3.4 C). Sphingolipids are well known for their roles in the immune response and shown to influence cell motility, signaling, and stimulation in a variety of immune cell types [35, 41, 42]. Their role in MSC immunomodulation is not clear, but several studies have looked at how bioactive sphingolipids affect over MSC immune function. One study saw that priming of MSCs with sphingosine-1-phosphate increased MSC mediated cardioprotection in a myocardial infarction mouse model [15] and reduced wall thickness and increased regeneration in a pulmonary artery hypertension in a rat model [34]. There is little evidence that sphingomyelinase activity within MSCs influences immunomodulatory outcomes. Here we propose a mechanism of action that not only explains morphological phenotypes related to potency, but may also influence overall MSC immune suppression and be used to tune MSC potency. Further studies are required to validate these findings.

## Conclusions

In this study, we characterized the morphological phenotype of four mesenchymal stem cell donors with differing IDO potency. We found that phenotypic morphological differences at normal cell culture conditions could discriminate between a high and low potency donor. We also found that PLSDA on four donors discriminated between donors according to potency and provided predictive power with 2.5% error rate by cross-validation method. A lipidomic profile assessment of 9 MSC donors binned into 4 high and 5 low IDO potency donors revealed key differences in lipid metabolism through a Pearson correlation heatmap. Further investigation, allowed us to identify that the production of long chain and complex ceramides was significantly higher in high IDO potency MSC donors versus low IDO potency MSC donors. We then saw that perturbation of the sphingolipid pathway in the production of ceramide by introducing exogenous bacterial derived neutral sphingomyelinase caused a large shift in morphological signatures in PLSDA. Using an IDO activity assay on neutral sphingomyelinase treated cells, we saw that there was a significant increase in both high and low IDO potency donors after treatment with neutral sphingomyelinase + interferon gamma when compared to interferon gamma treated MSCs. We establish that sphingolipids are a promising avenue for investigating the underlying mechanism of action for MSC immunomodulatory function that could also link the various morphological phenotypes that predict high and low potency donors.

## Notes

### Competing Interest Statement

The authors have declared no competing interest.

